# Global analysis of Two Component System (TCS) member of chickpea and other legume genomes implicates its role in enhanced nodulation

**DOI:** 10.1101/667741

**Authors:** Manish Tiwari, Manisha Yadav, Baljinder Singh, Vimal Pandey, Kashif Nawaz, Sabhyata Bhatia

## Abstract

Nodule organogenesis is governed primarily by phytohormone cytokinin. We observed the significant nodulation in chickpea at particular cytokinin concentration (2.5×10^−7^) which indicated the importance of cytokinin in nodule development. Cytokinin signaling is mediated through the <u>T</u>wo <u>C</u>omponent <u>S</u>ystem (TCS) which comprises of sensor histidine kinases (HKs), histidine phosphotransfer proteins (HPs), and response regulators (RRs). Therefore, we analyzed the interconnection of cytokinin with TCS molecules during root nodule development through global analysis of TCS candidates in legumes with special consideration to cytokinin receptor and Type-B RR member. We have conducted an in depth global analysis of TCS family members in chickpea and other legumes, *Medicago* and pigeon pea. Higher number of TCS genes were found in *Medicago* (96), followed by pigeonpea (75) and chickpea (67). A good correlation between TCS members with their corresponding total number of genes were observed in all three-legume species. Collinearity analysis of TCS revealed phylogenetically closer proximity of *Cicer* to *Medicago* followed by *Glycine* than *Cajanus*. Comprehensive analysis of 3-dimensional structure, genomic organisation and domain arrangement showed a conservation of TCS members within species. In depth investigation showed that HKs were mainly conserved among TCS members in legumes and non-legumes while divergence occurred at level of RRs. Further, Type-B RRs were functionally most diversified in RRs based on phylogeny, syntenic and transcript analysis. Few numbers of segmentally duplicated pair of TCS showed difference in their transcriptional regulation suggesting the functional evolution. For functional characterization the *cre1* mutants of (*Medicago*) were complemented with chickpea cytokinin responsive HKs and nodulation deficient phenotype of mutants were restored. A synchronous cytokinin-induced expression of chickpea cytokinin receptor *HKs* and *CaNIN* provides strong relation of cytokinin signaling during nodulation. Furthermore, interesting potential candidate *CaRR13* was selected to deduce the underlying molecular mechanism of nodulation, chickpea in specific and legumes in general.

## Introduction

Cytokinin plays a major role in promoting apical dominance, shoot-root branching, seed germination, leaf senescence, floral transition, photosynthesis and lateral bud growth (Werner et al., 2001). Ectopic cytokinin treatment leads to the formation of nodule-like structure even in the absence of rhizobia. Besides nodule formation and maintenance, it also controls the number of root nodules formed through shoot derived inhibitors. Even a gain in function mutation of LHK1, a CRE1 ortholog of Arabidopsis has been reported to result in spontaneous nodule formation, while loss of function resulted in impaired nodulation (Murray et al., 2007; Tirichine et al., 2007). Cytokinin signal transduction is mediated by two component system (To and Kieber, 2008). The presence of TCS is reported in all form of life such as eubacteria, archaea and eukaryotic organisms except animalia kingdom. They are the key regulatory molecules which control many biological processes, including cell division, proliferation, growth and stress (Pareek et al., 2006; MIZUNO, 2005; Chen et al., 2012). TCS or better known as His-to-Asp phosphorelay consist of a sensory receptor, i.e. histidine kinase, which possess His-kinase (HK) domain and a receiver (Rec) domain, a phosphotransfer protein histidine phosphotransferase (HP) and effectors, response regulator (RR). In response to the stimulus, kinase domain of HK auto phosphorylates itself at histidine amino acid residue. Further, the signal is transferred downstream via phosphoryl transfer through HP to the aspartic acid residue of Rec domain of a Type-B RR (West and Stock, 2001). Type-B RR can act as transcription factor through binding cis-elements in promoter region of the target genes such as NSP2 and bLHLH.

Histidine kinases are classified into various subclasses on the basis of their domain organization such as cytokinin receptor (CHASE, cyclase/HK-associated sensory extracellular) (Alderete et al., 2001), ethylene receptor (ethylene binding domain) and phytochrome receptor (PHY and PAS). Apart from these HK, *Arabidopsis* cytokinin insensitive1 (CKI1), *Arabidopsis* histidine kinase1 (AHK1) and *Arabidopsis* histidine kinase5 (AHK5) are also annotated in the TCS family. A highly conserved xHQxKGSSxS motif of AHPs mediates a phosphate transfer from Rec domain of HKs to the Rec domain of RRs. Conserved amino acid residue histidine is required for phosphate group transfer and hence downstream TCS signaling. This conserved histidine is absent in few of histidine phosphotransferase protein, considered as pseudo histidine phosphotransferase (PHP). PHP functions as a negative regulator of cytokinin signaling (El-Showk et al., 2013). Response regulators can be broadly categorized into Type-A, Type-B, Type-C and PseudoRRs. Type-A and Type-C RRs contain REC and short C-terminal extension, and both act as negative regulators of cytokinin signaling. Type-A RRs are induced by cytokinin treatment while there is no effect on Type-C RRs expression (To et al., 2007, 2004; To and Kieber, 2008). A distinguishing feature is present in the C-terminals of Clock PRRs; Co, Col and Toc1 motif and functions in the regulation of circadian rhythms(Mizuno and Nakamichi, 2005; Makino et al., 2000; Nakamichi et al., 2010). Type-B RRs contain a REC domain at N terminal followed by a large domain, GARP (GOLDEN/ARR/Psr1, ~60 amino acids motif) at the C terminal which have a distant similarity with Myb DNA binding superfamily (Tajima et al., 2004; Mason et al., 2004). Additionally, there is a diverged group of RRs which lack the conserved aspartate residue required for phosphorylation and termed as pseudo RRs.

Two component system genes were implicated in stress condition, and show an enhanced expression (Fujita et al., 2005). Already TCS genes were explored at the whole genome level in various plant species such as *Arabidopsis thaliana, Lotus japonicus, Physcomitrella patens*, soybean, maize, rice and Chinese cabbage (Liu et al., 2014). All investigations so far provided enough evidence for their genic and structural organization of TCS family, however an important aspect regarding their evolution, diversification and transcriptional regulation in legumes lineage and its involvement in nodulation is still unexplored. There is a need to conduct a study for systematic investigation of TCS evolution in legumes for their functional diversification and connection to nodulation.

Legumes especially chickpea and pigeonpea are widely cultivated under diverse conditions in India. In the present investigation, we have selected these crops along with model crop *Medicago* to analyze global exploration of TCS in legume lineage. A vast study on whole genome was performed to identify TCS genes and their associated properties such as, the genomic and structured organization in legumes. Integration of phylogeny, transcriptional regulation and evolutionary analysis of TCS genes provide useful insight of their functional diversification. Evolutionary aspect pertaining to divergence time of legumes in reference to each other and to non-legumes and their specification in legumes has also been deduced. Chickpea HKs were functionally validated for cytokinin receptor activity by complementation in yeast and *Medicago* mutants. Further, member of Type-B RR was selected based on its transcriptional regulation and evolutionary importance to provide an insight into its *in planta* role during root nodulation, chickpea in particular and legume species in general. Our understanding of key elements of TCS family would provide insights about the cytokinin induced TCS signaling which can be used to improve nodulation phenotype. This can be used to facilitate the targeted manipulation of metabolic pathways in order to increase plant productivity, stress-adaptation and quality improvement in chickpea.

## Materials and Methods

### Identification of TCS genes in chickpea, *Medicago* and pigeonpea

Protein sequence from chickpea, *Medicago* and pigeonpea were downloaded from an online database of the respective species. The proteome database of kabuli was downloaded from the FTP server of NCBI (ftp://ftp.ncbi.nlm.nih.gov/genomes/Cicer_arietinum/) while the desi has been downloaded from NIPGR Chickpea Database (http://nipgr.res.in/CGAP2/download/genome_sequencing/annotation/Gene%20annotation/CGAP_v2.0/Ca_Pep_v2.0.fa). The *M. truncatula* proteome database was downloaded from http://www.Medicagogenome.oru/downloads and *C. cajan* from (http://gigadb.org/dataset/100028). TCS protein sequences from *Arabidopsis thaliana, Oryza sativa, Lotus japonicus, Glycine max* and *Brassica rapa* were downloaded and used as query to perform BLAST. BLASTP and HMMER was used to extract TCS sequences in all three legumes. All the sequences having an e-value of 10 were extracted and subjected to SMART (http://smart.embl-heidelberg.de/), CDD (http://www.ncbi.nlm.nih.gov/Structure/cdd/wrpsb.cgi) and Pfam (http://pfam.janelia.org/) database for domain analysis.

### Structural information, motif identification, subcellular localization and phylogenetic analysis

The exonic-intronic information was visualized using Genetic Structure Display Server (http://gsds.cbi.pku.edu.cn/). Protein modelling was performed using Phyre^2^ (Kelley et al., 2015) and conserved motifs were identified using MEME (Multiple Expectation Maximization for Motif Elicitation, http://meme.nbcr.net/meme3/meme.html). Parameters used were distribution of motif occurrences: zero or one per sequence, maximum number of motifs: 25 and optimum motif width: ≥ 6 and ≤ 50. Functional annotation and subcellular localization of TCS genes were done using Blast2GO (Conesa et al., 2005). The multiple alignment of protein sequence of all the aforementioned organisms was performed by MUSCLE program keeping gap penalty −2.9. The unrooted tree was constructed for phylogenetic analysis via Neighbour-Joining (NJ) method keeping the bootstrap value 1000 in MEGA 6.0 (http://www.megasoftware.net/) and visualized through iTOL v 3(http://itol.embl.de/).

### Chromosomal localization and evolutionary analysis

The genomic position of chickpea, *Medicago* and pigeonpea provided in the LIS database (http://legumeinfo.org/) was used to map the positions of TCS family through MapChart (https://www.wageningenur.nl/en/show/Mapchart.htm) (Voorrips, 1994). The estimation of segmental pairs and synteny analysis was carried out using data from Plant Genome Duplication Database. Time of divergence was calculated using the synonymous mutation rate of substitutions per synonymous site per year as per chalcone synthase gene where, T= Ks/1.5X10^−8^ (Koch et al., 2000)

### Digital expression analysis of TCS genes in chickpea, *Medicago* and pigeonpea

Tissue specific transcriptomes of chickpea for various tissues and stresses such as leaf (SRX048833), root (SRX048832), flower bud (SRX048834), pod (SRX048835), seed (SRX125162) and nodule (PRJNA214031) were downloaded from SRA (Sequence Read Archive. Stress-specific (salt, desiccation and cold) transcriptome of chickpea root and shoot were retrieved from the SRA database (SRP034839) (Garg et al., 2015). Reads were filtered, trimmed and mapped on the chickpea (desi) genome using TopHat2 (Kim et al., 2013). Cuffdiff was used to determine the differential gene expression value in FPKM and heat maps were visualized through MeV software.

### Plant growth, treatments, and sample harvesting

Chickpea seeds were surface sterilized, kept on the germination sheet for two days and then transferred to pot containing vermiculite and peat (3:1) under environmentally control conditions. Root, leaves, seeds and flowers were harvested and used for tissue or organ-specific expression analysis. For nodule-specific expression, young seedlings were infected with *M. ciceri* and transferred to the sand in Nitrogen free McKnight’s solution for 10 days at 22 ± 1 °C with 16:8-h photoperiod. For hormone treatment, 100 μM 6-BA (6-Benzylaminopurine) of cytokinin and 100 μM + (−) Abscisic acid, were sprayed to foliar region. Root and shoots were collected at 1, 3, 5h and control plants were sprayed with double distilled water. For stress treatment, plants were uprooted carefully and kept on the folds of tissue paper to study dehydration effect. To study the effects of cold and salt stress, uprooted plants were shifted to 4°C and 150 mM NaCl in the 500ml water respectively. All the treatments were done for 1, 3 and 5 h while the control plants were kept in similar conditions with 500ml of distilled water.

### RNA extraction and quantitative reverse transcription PCR analysis

RNA was extracted using LiCl method (Choudhary et al., 2009) and quality was checked on Bio-analyser. First strand cDNA was synthesized using Bio-Rad iScript cDNA synthesis kit by following the manufacturer’s protocol. Primers were designed using Primer Express software v2.0 and qRT-PCR was performed using EF1α as an endogenous control. Three biological and technical replicates were kept for each experiment and ΔΔCT method was used to calculate the relative expression

## Results

### Identification and annotation of TCS family in legumes

In the present investigation, we identified 67 TCS members in chickpea (desi and kabuli), 96 in *Medicago* and 75 in pigeonpea. Few genes were common in both the chickpea varieties while some were present exclusively in kabuli (**Figure 1 and Supplementary table S1**). There are three main components of TCS *viz*., HKs (19 in chickpea, 21 in *Medicago* and 20 in pigeonpea), HPs (6 in chickpea, 8 in *Medicago* and 8 in pigeonpea), and RRs (40 in chickpea, 58 in *Medicago* and 39 in pigeonpea). Domain and motif alignment analysis revealed that HK in kinases, HPt in phospho-transferases, rec and Myb in Type-B RRs and rec in other RRs were very well conserved. Despite of these conserved domains, we examined that there was a class of diverged two-component elements, namely, HKLs (25 in *Medicago* and 30 in pigeonpea) (**Supplementary Table S2**), PHPs (2 in chickpea, 2 in *Medicago* and 2 in pigeonpea), and pseudo-RRs (6 in chickpea, 9 in *Medicago* and 6 in pigeonpea) were devoid of conserved phosphorylation sites. Additionally, RRs are subclassified in Type-A (9 in chickpea, 11 in *Medicago* and 12 in pigeonpea), Type-B (16 in chickpea, 35 in *Medicago* and 17 in pigeonpea) and Type-C (5 in chickpea, 12 in *Medicago* and 10 in pigeonpea) (**Figure 1 and Supplementary table S1**). Additionally, the ratio of the total number of genes in *Medicago* as compared to chickpea and pigeonpea was 1.5 and 1.03, respectively and pigeonpea to chickpea was 1.4. Similarly, the ratio of TCS genes between Medicago-chickpea, *Medicago*-pigeonpea and pigeonpea-chickpea was 1.43, 1.28 and 1.11, respectively (**Supplementary Table S9**).

**Figure 1.**
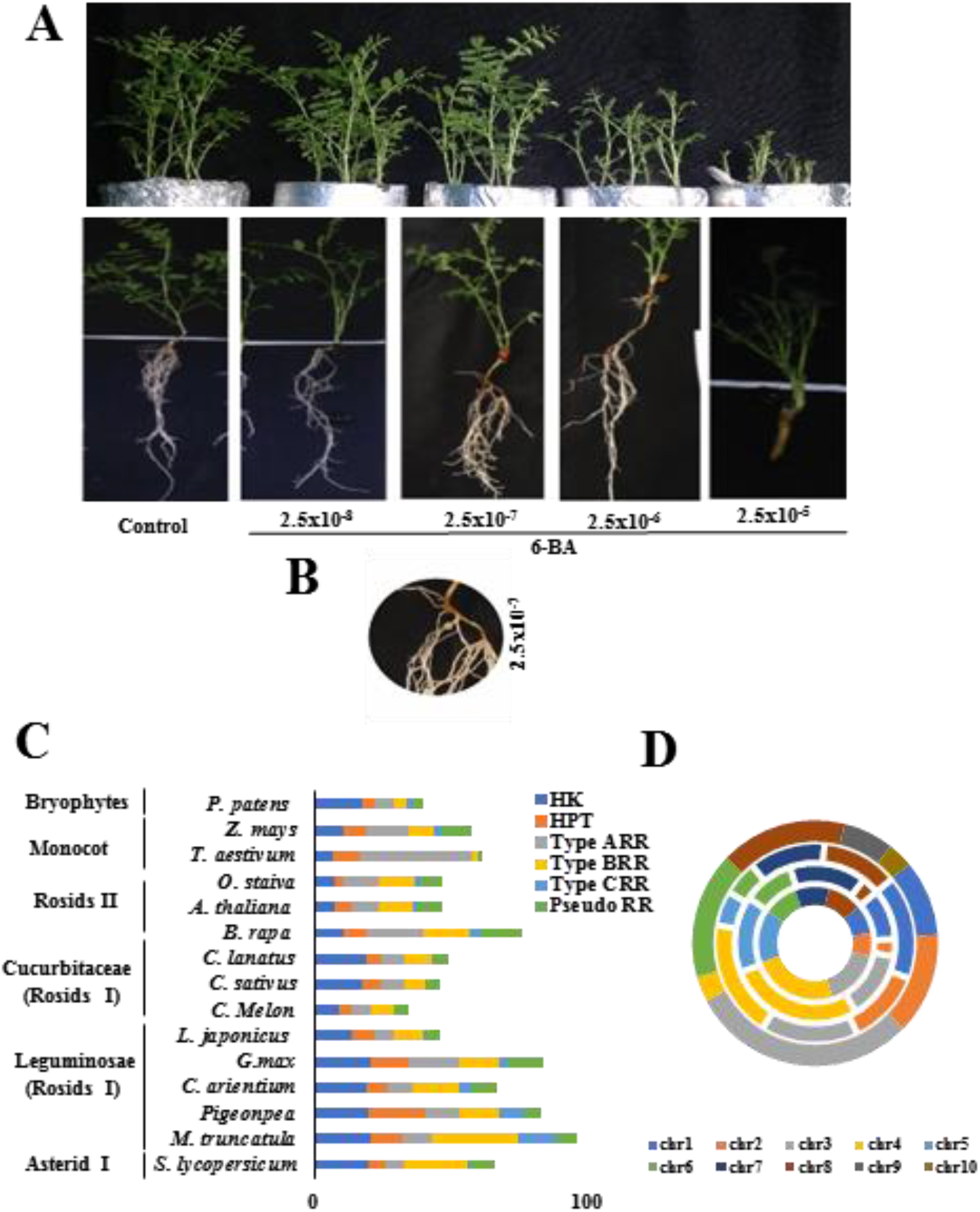
Interconnection between cytokinin hormone and TCS. **A.** Images represent the shoot and root phenotype of chickpea under varying concentration of cytokinin. **B.** Induction of root nodule at 2.5×10^−7^ 6-BA. The experiments were carried in triplicates. **C.** The distribution of HK, HP, Type-A RR, Type-B RR, Type-C RR and Pseudo RRs in various crop species. **D.** Doughnut denotes the frequency of TCS genes in the chromosome of cv. desi chickpea (inner circle), cv. kabuli chickpea, barrel medic and pigeonpea (outer circle).

### Integrative analysis of phylogeny, synteny and transcriptional regulation of TCS among legumes

The HK(L) were further sub grouped in six families such as cytokinin receptors, ethylene receptors, AHK5/CKI2, AHK1, CKI1 and phytochromes. Four *Ca*HKs (*Ca*HK7,14,18,19), five *Mt*HKs (*Mt*HK11, 12,16, 20, 21) and four *Cc*HKs (*Cc*Hk1,5,7,13) represents cytokinin receptor subfamily (**Figure 2**). The paralog pairs of cytokinin receptors were also orthologous pairs between chickpea and *Medicago* (*Ca*HK7-*Ca*HK18, *Mt*HK12-*MtH*K20, *Ca*HK7-*Mt*HK20 and *Ca*HK18-*Mt*HK12) implicating their importance in cytokinin signaling (**Supplementary Table S4, S5 and S6**). Similar tissue-specific expression was observed in orthologous pairs (*Ca*HK7-*Mt*HK20 and *Ca*HK18-*Mt*HK12) while paralogs (*Ca*HK7-*Ca*HK18 and *Mt*HK12-*MtHK*20) showed antagonistic expression. Similarly, paralog pairs of cytokinin insensitive family were orthologous pairs in chickpea and *Medicago* (*Ca*HK4-*Ca*HK8, *Mt*HK1-*MtHK*13, *Ca*HK8-*Mt*HK1 and *Ca*HK4-*Mt*HK13). The digital expression of orthologous and paralogous pairs was similar depicting no functional diversification in members of the cytokinin insensitive family (**Supplementary Table S4, S5 and S6**). Generally, members of cytokinin insensitive family were found to be expressing. However, CcHK3, MtHK15 and CaHK3 showed induction in expression specifically during nodulation. Additionally, ethylene receptor subfamily (I and II) and phytochrome subfamily was also explored in three legumes (**Figure 2 and Supplementary Table S1**). The paralogs of ethylene receptor family are also orthologs among chickpea, *Medicago* and pigeonpea (**Supplementary Table S6 and S5**).

**Figure 2.**
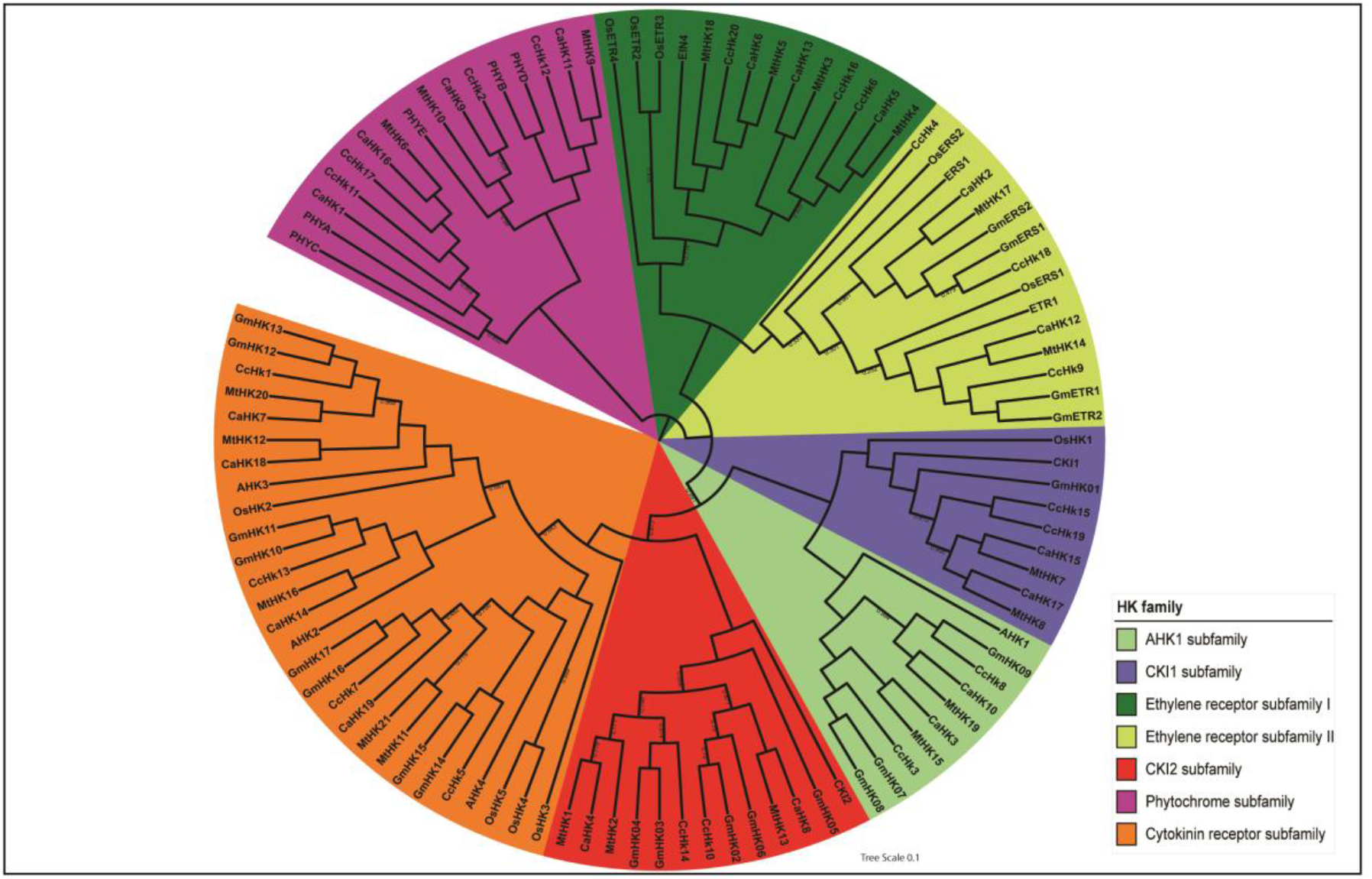
Phylogram of histidine kinase. The phylogenetic analysis classified the histidine kinase family into ethylene receptor, cytokinin receptor, phytochrome, AHK1, CKI and CKI2. The bootstrap values are depicted as numbers on clades.

An interesting observation on phylogram of HPs was that all the members of *Oryza sativa* fall in a distinct clade. Cytokinin signaling positive regulator AHP1, showed a closer relationship with *Ca*HP1, *Mt*HP3 and *Cc*HP5-6, and AHP2, 3, 5 with *Ca*HP5, *Mt*HP2, *Cc*HP2. The members of cytokinin positive regulators were segmentally duplicated within as well as between chickpea and *Medicago* (**Supplementary Table S4**). Similarly, cytokinin signaling negative regulator AHP4 showed a strong relationship with CaHP4, CaHP6, MtHP4-5 and CcHP3-4, (**Figure 3**) however only orthologous pairs were found in chickpea and *Medicago* but paralogs were absent. These analyses dictate that cytokinin positive regulators were more diverged as compared to the negative regulators (Supplementary Table S4).

**Figure 3.**
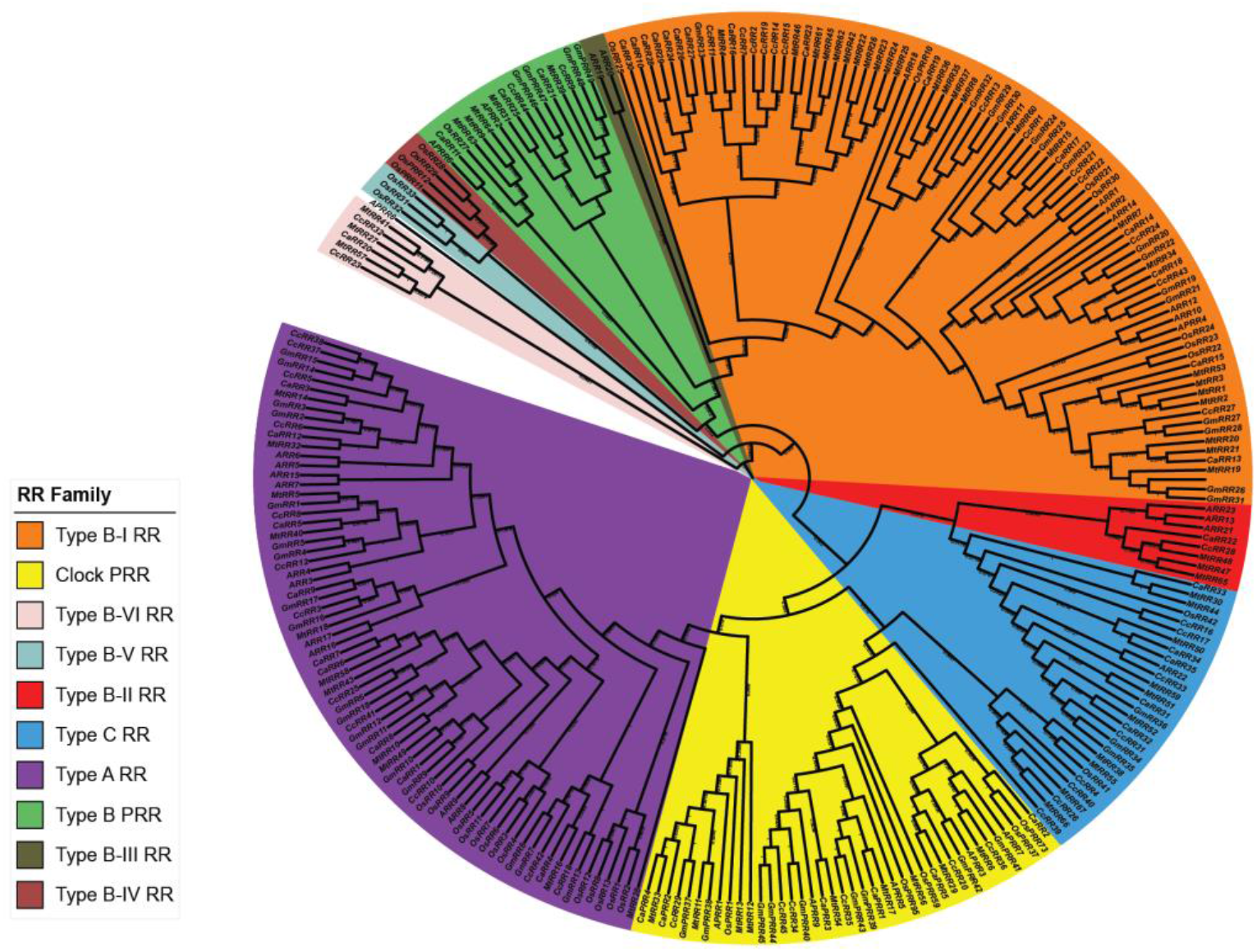
Phylogram of response regulator. The phylogenetic tree classified the response regulator family into TYPE-A RR, TYPE-B RR, TYPE-C RR and Pseudo-RR subfamily. The bootstrap values are depicted as numbers on clades.

Type-A RRs paralogs numbered 3 in *Medicago*, 2 in pigeonpea and 1 in chickpea. Similarly, the number of Type-B RR paralogs is 5 in *Medicago*, 1 in pigeonpea and 2 in chickpea (**Supplementary Table S6**). Four ortholog pair of Type-A RR and eight ortholog pairs of Type-BRRs had paralogs in chickpea and *Medicago*. However, no paralogs of Type-C RRs were found in any of the three-legume species (**Supplementary Table S6**). This result indicated that Type-B RRs were comparatively more diverged among the RRs. Type-B RRs are very complex and further classified into six subgroups (I-VI) unlike Type-A RRs and C RRs (**Figure 3**). Among six subgroups, members of Type-B RRs of chickpea, *Medicago* and pigeonpea clustered with other Type-B RRs in phylogram and sub grouped as Type-B I RRs and II RRs. Interestingly, we observed a legume-specific cluster of Type-B RRs (*Ca*RR20, *Mt*RR27, 41, 57 and *Cc*RR23, 32) in phylogram and assigned as Type-B VI RRs (**Figure 3**). However, Type-B IV and V RRs contains only rice proteins indicating the divergence of monocots from dicot species. Two orthologous pairs of Type C-RRs were found in chickpea-*Medicago*, chickpea-pigeonpea and pigeonpea-*Medicago* (**Supplementary Table S4 and S5**). Additionally, diverged TCS Pseudo-RRs were divided into Clock PRR and Type-B PRR. Intriguingly, *Ca*RR2, Type A-RR, falls in clock PRR’s clade in the phylogenetic tree due to the presence of same motif as found in *Ca*PRR5 (**Figure 3**).

### Collinearity analysis depicted the rearrangement of TCS members on chromosome among legumes

In all TCS members, 57, 91 and 41 were localized on chickpea, *Medicago* and pigeonpea chromosomes respectively (**Supplementary Figure S4**). The maximum TCS segmental pairs of Ca-Mt (chromosome of chickpea-chromosome of *Medicago*) were found on the chromosome Ca4-Mt1 (12 pairs), Ca5-Mt3 (9 pairs) and Ca5-Mt3 (6 pairs) which corroborates the previous published syntenic relationship between chickpea and *Medicago* genome (**Supplementary Table S7**). Collinearity analysis of chickpea TCS with other legumes revealed that the higher number of segmental pairs were found on chromosome number 4 of chickpea as compared to soybean and *Medicago* (Ca4-Mt1, Ca4-Gm10, Ca4-Gm20 and Ca4-Gm3) (**Supplementary Table S7**). Similarly, collinearity analyses of *Medicago* TCS showed that the higher number of segmental pairs were found on chromosome number 1 of *Medicago* with respect to soybean and chickpea (Mt1-Ca4, Mt1-Gm10, Mt1-Gm20 and Mt1-Gm3) (**Supplementary Table S7**). In pigeonpea, maximum number of orthologs were present on chromosome number 3, however no specific synteny with any chromosome of other legumes was observed (**Supplementary Table S7**). This analysis indicated that there is a common ancestry between legumes and depicts a close relationship of chickpea with *Medicago* and soybean.

### Divergence of TCS members between legumes and non-legume species

In depth analysis of the pattern of evolution and divergence of TCSs among legumes and with non-legume species was performed. Relative Ks value was used to calculate the divergence time of TCS in *Cicer, Medicago, Cajanus, Glycine* and *Arabidopsis*. The duplicated orthologous gene pair of chickpea, pigeonpea and *Medicago* with *Arabidopsis* showed a peak at Ks value of 2.1-2.2 which corresponds to ~138 to 146 Mya. Similarly, the relative Ks value of 0.3-0.4, 0.5-0.6 and 0.7-0.8 for orthologs between chickpea-Medicago, chickpea-soybean and chickpea-pigeonpea corresponds to a divergence time of 20-27, 33-40 and 47-53 Mya respectively (**Figure 4**). This data is in accordance with previous findings that chickpea is phylogenetically closer to *Medicago*, followed by soybean, pigeonpea and *Arabidopsis*.

**Figure 4.**
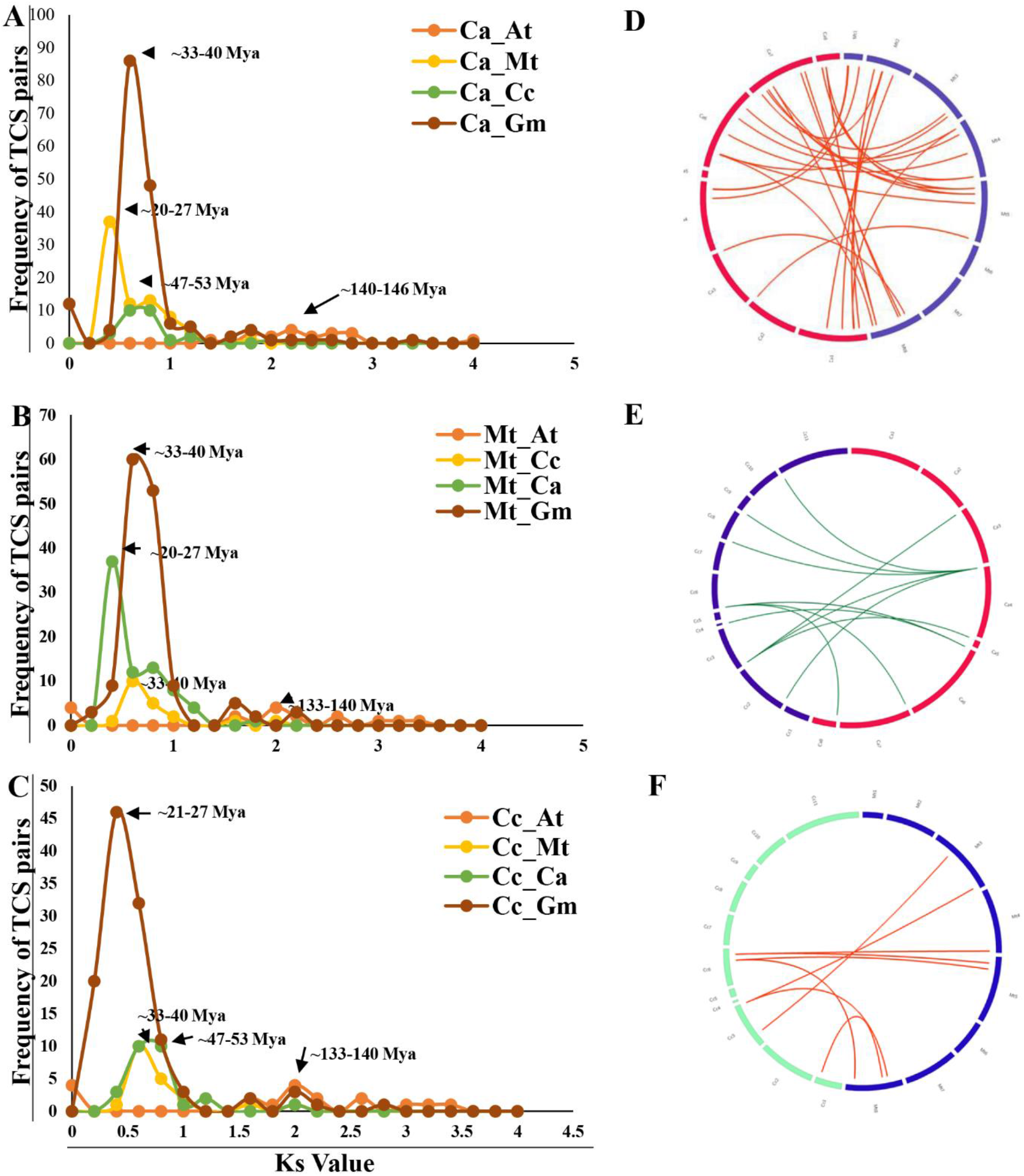
Synteny and collinearity analysis of TCS members among legumes. **A-C.** Graph represents Ks values and number of segmentally duplicated pairs indicating divergence time among the legumes and non-legumes. The Y-axis indicates the numbers of segmentally duplicated pairs and the horizontal axis indicates the Ks values with a 0.2 interval. **D-F.** Circos showed the number and position of orthologous pairs in chickpea-*Medicago*, chickpea-pigeonpea and *Medicago*-pigeonpea.

Genome-wide median value for the Ka/Ks ratio is 0.09 to 0.18 for HKs and 0.23 to 0.37 for RRs orthologs between different legumes along with *Arabidopsis*. The lower value of Ka/Ks for HKs indicated that strong purifying selection acted during evolution while the higher Ka/Ks ratio of RRs indicated an increase in nonsynonymous substitution as compared to synonymous substitution (**Figure 5D and 5E**) This was also validated by analyzing the mean value of the Ka ratio in all the segmentally duplicated HK and RR pairs were found to be ~2, whereas the mean value of Ks ratio was ~1 (p-value-0.00001). The higher Ka/Ks ratio of RRs denoted the recent positive selection among TCS components (**Supplementary Table 4 and 5**).

**Figure 5.**
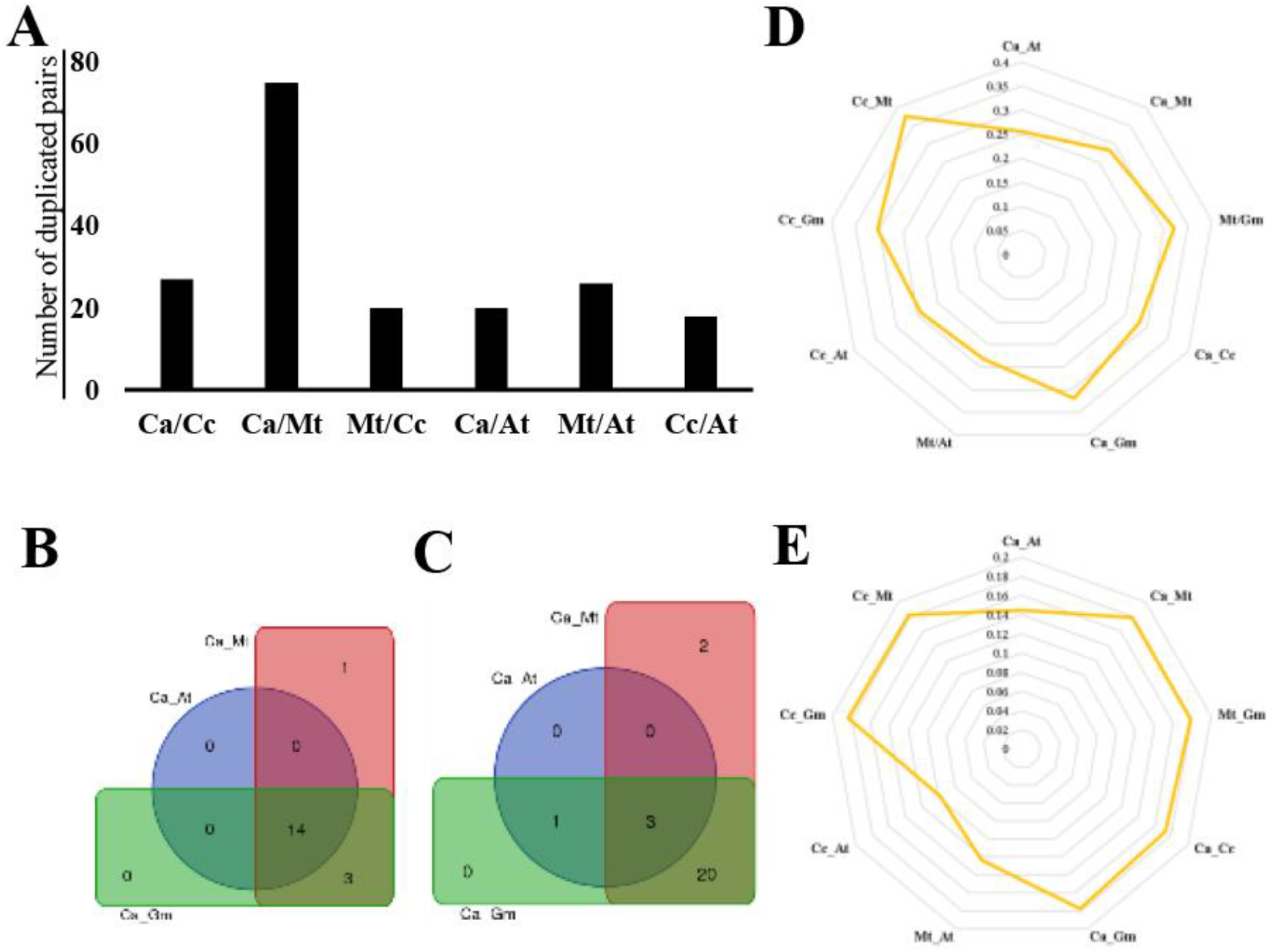
Evolutionary analysis of TCS members. **A.** Bar diagram represents the pair-wise segmentally duplicated TCS members in plants. **B-C.** Venn diagram denotes the pair-wise comparison number of HKs **(B)** and RRs **(C)** in plants. **D-E.** Spider web indicates pair-wise comparison of genome wide median value of Ka/Ks ratio of HKs **(D)** and RRs **(E)** in plants.

To explore whether the existence of paralogs may or may not instigate the evolution of protein-coding genes, we examined the conservation of gene orthologs in absence or presence of paralogs. Through our analysis between *Medicago* and chickpea we found that orthologs with paralogs did not behave differently from genes without paralogs. Comparative analysis of duplicated pairs between Ca_At, Ca_Gm and Ca_Mt revealed that out of 19 chickpea HKs, 14 were duplicated with Arabidopsis, 17 with soybean and 18 with *Medicago* and that too with purifying selection (lower Ka/Ks value) (**Figure 5B, Supplementary Table S3 and S4**). Whereas only 4 RRs out of 40 in chickpea were found to be duplicated in comparison to Arabidopsis. Surprisingly, the number rose significantly to 24 and 25 in soybean and *Medicago*, respectively. The results clearly indicated a legume specific duplication and diversification of RRs (**Figure 5C**).

### *In silico* annotation of TCS revealed the difference in their functional role

To analyze the functional divergence among legumes, we extracted the relative expression values of different tissues. Most of segmentally duplicated pairs between legumes showed that expression of orthologs was conserved. However, we also found that few of the TCS orthologs (28 in chickpea-*Medicago*, 7 in chickpea-pigeonpea and 4 in pigeonpea-*Medicago* showed opposite tissue-wise expression indicating the diverged functional role in TCS across the legumes (**Figure 6A, 6B and 6C**). Digital expression of ortholog pairs (*Ca*RR9-*Cc*RR3 and *Ca*HK4-*Cc*HK14) of chickpea and pigeonpea showed contrasting expression in nodule tissue (**Figure 6A, 6C and Supplementary Table 4**).

**Figure 6.**
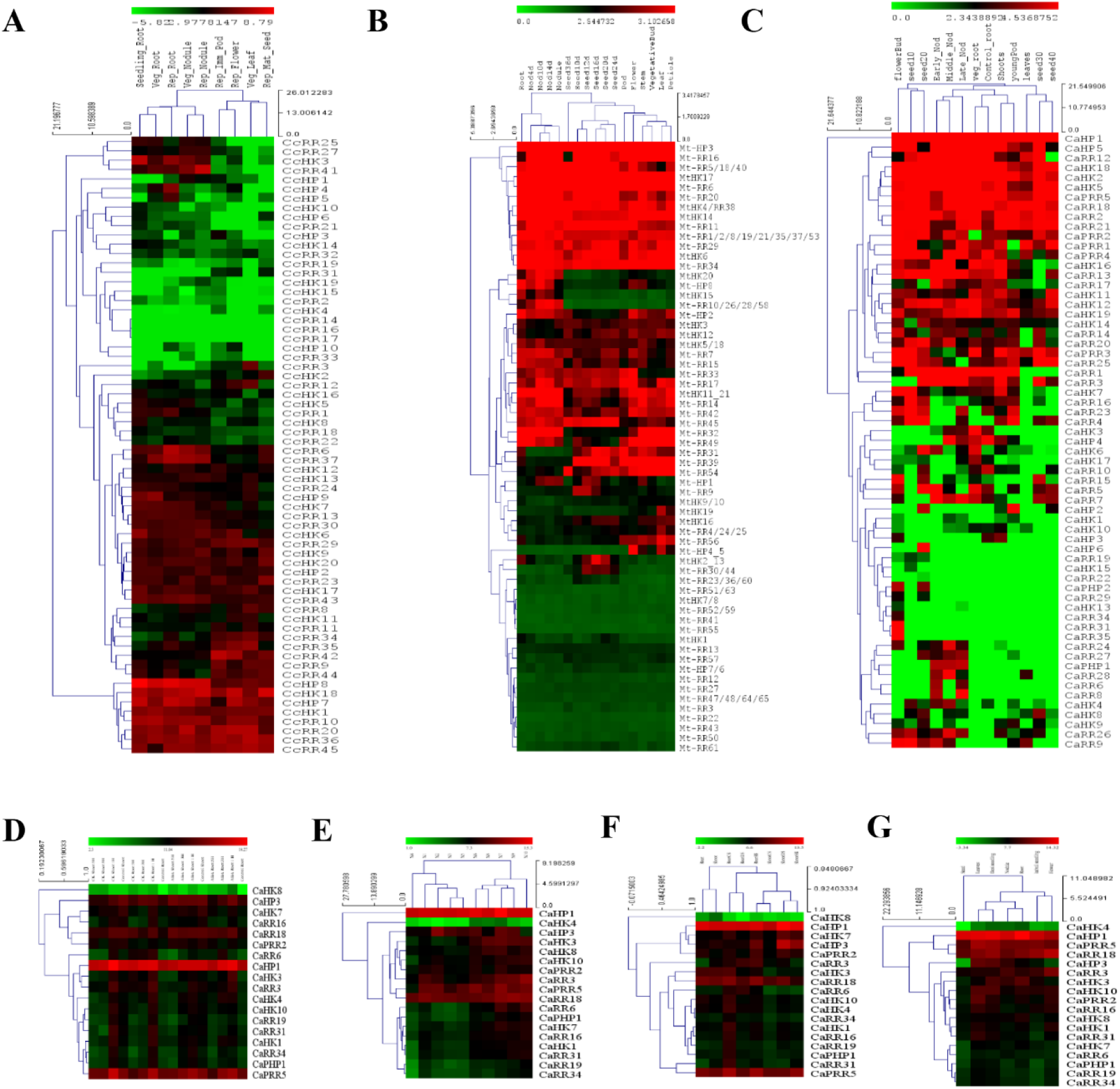
Spatiotemporal and stress-responsive expression analyses of TCS members. **A-C.** The digital expression analysis in different tissues of pigeonpea **(A),** *Medicago* **(B)** and chickpea **(C). D-G.** Heat map denotes the expression analysis of TCS members during cytokinin-ABA treatment in root and shoot **(D),** Nodule-specific stages (N1-1HPI, N2-3HPI, N3-6HPI, N4-12HPI, N5-24HPI, N6-3DPI, N7-7DPI, N8-14DPI, N9-21DPI, N10-28DPI) (E), Abiotic stress in root and shoot **(F)** and different tissues **(G).** The experiments were carried in triplicates and expressed as means ± S.E

*In silico* functional annotation exposed that members of *Mt*TCS and *Cc*TCS were localized to the different compartment to perform different functions *viz*., reproduction, lipid metabolism, cell death and embryo development. However, members of *Ca*TCS were solely dedicated to signal transduction *via* the formation of macromolecular complex (**Supplementary Figure S3**).

### Role of TCS genes of chickpea in various developmental and stress condition

Eighteen out of 67 *Ca*TCS genes were selected on the basis of significant expression using the available data at CTDB, nodule and seed transcriptome (Pradhan et al., 2014) to perform the comprehensive expression analysis in various tissues using qRT-PCR. TCS genes *viz*., *Ca*HK3, *Ca*HK4, *Ca*HK7, *Ca*HP1, *Ca*PRR5, *Ca*RR3 and *Ca*RR18 were found to be highly expressed in roots. The higher expression of *Ca*HK7 indicated its active participation in cytokinin signalling in roots. Since TCS genes played a crucial role in nodule formation, therefore we also analysed the expression in nodule insilico and through qRT-PCR expression analysis. We observed a set of TCS genes such as *Ca*HP1, *Ca*HP3, *Ca*PRR5, *Ca*RR3, *Ca*RR13 and *Ca*RR18 were active throughout nodulation (**Figure 6F**). The expression of TCS genes was induced onwards 24 hpi and maintained till 3 weeks. However, expression of RR members was higher as compared to HK members in mature nodules. The higher expression of TCS genes in seedlings of desi cultivar as compared to kabuli cultivar, indicated cultivar-specific responses (**Figure 6G**). Besides these functions, *Ca*TCS genes also played a very important role in the development of flower and leaf, and in different abiotic stress conditions (**Figure 6B, 6F and 6G**).

Previous studies based on qRT-PCR, Northern blotting and microarray has shown that there is no effect of exogenous cytokinin or ABA on HK, HP and Type-B RR genes in *Arabidopsis*. However, major response to cytokinin treatment is shown by Type-A RR (Brenner et al., 2005; Ha et al., 2012; Rashotte et al., 2011). Therefore, to determine the effect of exogenous cytokinin or ABA on the TCS gene expression in chickpea, hormone spraying on the foliar region was performed. Interestingly, the induced expression of *Ca*TCS genes (except *Ca*HK8) was observed 1 h and 3h post cytokinin treatment as compared to 5h (**Figure 6D**). Cytokinin works antafonistically to ABA during most stress conditions. (Murray et al., 2007; Tran et al., 2010). The suppression of *Ca*TCS genes was observed in shoot, while elevated expression observed in roots. ABA perception is first achieved in root tissue which passes the signal to shoot tissue and participate in regulation of various physiological activities (Urao et al., 1998; Hirayama and Shinozaki, 2007).

### Role of chickpea cytokinin receptors and Type-B RRs in root nodulation

To further establish the role of HKs during cytokinin signaling and nodulation, expression analysis was performed at different concentrations of cytokinin (2.5*10^−6^ and 2.5*10^−7^) along with control. The expression of HKs started elevating at 12 h post treatment and much higher elevation was observed upon treatment with higher cytokinin concentration. Among the cytokinin receptor, CaHK19 showed exceptional response towards cytokinin. Intriguingly, NIN was also found to be elevated along with HKs at 12 h post treatment (**Figure 7A**). The results indicated a link between cytokinin reception and nodulation downstream signaling. To functionally validate the *CaHKs* as true cytokinin receptors, complementation of HKs in *Medicago* mutant, *cre1* along with yeast mutant, *sln1Δ* was employed. All four cytokinin receptors were able to restore the nodule deficient phenotype of *cre1* mutants upon complementation through hairy root transformation (**Figure 7C**). Similarly, in *S. cerevisiae* complemented yeast mutants (*Δsln1: Ca*HK14 and *Δsln1: Ca*HK18) revived efficiently in cytokinin-dependent manner as compared to mutant and vector control (**Figure 7B**).

**Figure 7.**
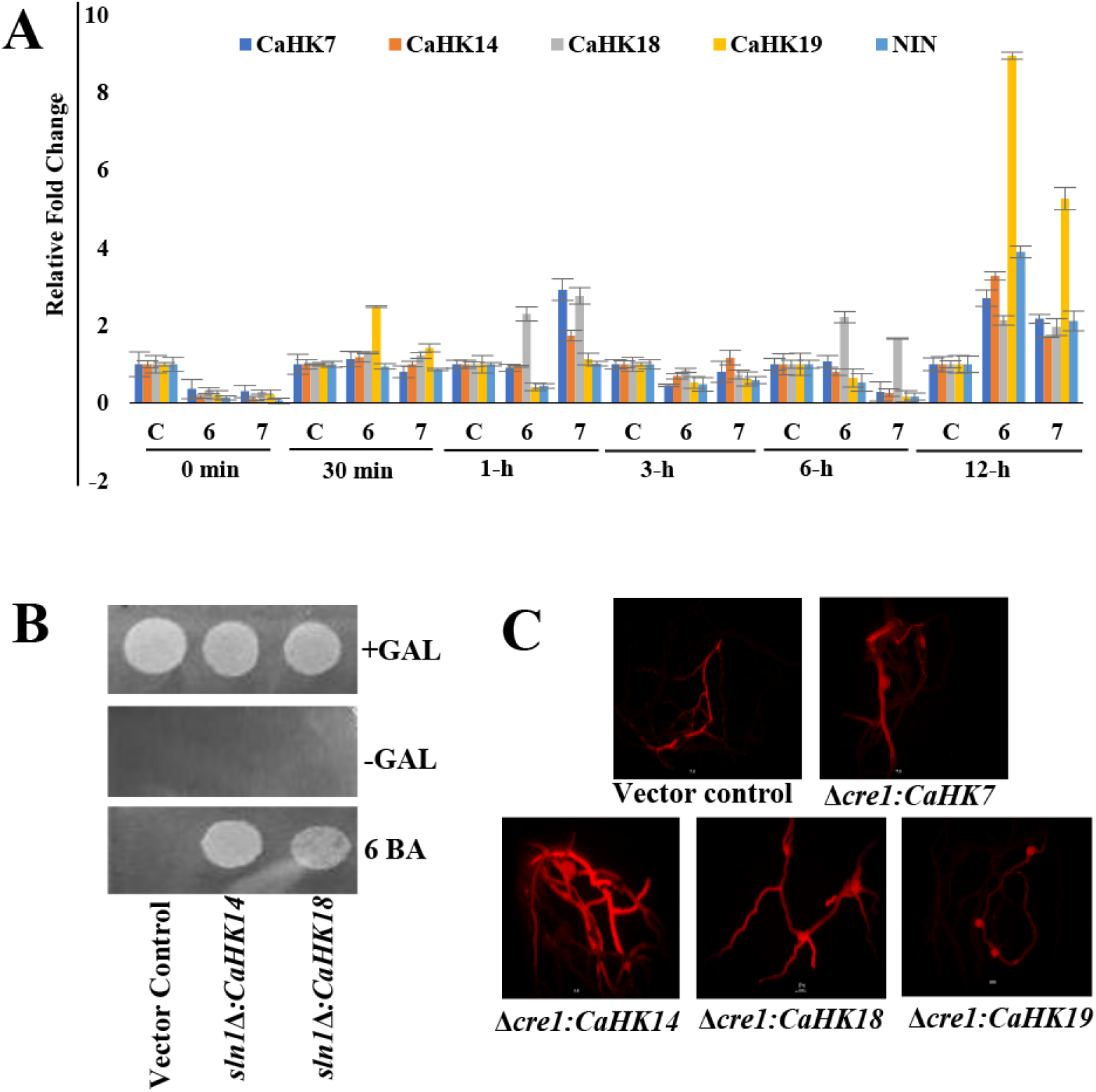
Expression and heterologous expression of HKs. **A.** Bar diagram represents the expression profile of *Ca*HKs during control (C) and 6BA (6 denotes 2.5X10^−6^ M and 7 denotes 2.5×10^−7^ M) at 30min, 1h, 3h, 6h and 12h. The experiments were carried in triplicates and expressed as means ± S.E.**B.** Complementation of yeast mutant sln1Δ with CaHK14 and CaHK18. **C.** Complementation of *Δcre1* mutant with CaHKs in *Medicago*

As per evolutionary analysis and also previous investigation, it was evident that RRs diverged functionally while HKs remain conserved. Type-B RRs were more variable among the RRs and also act as a transcription factor. Type-B RRs, *CaRR13* was selected based on its continuous elevated expression, especially in nodule during cytokinin treatment as well as *M.ciceri* infection (**Figure 8A**). Orthologs of CaRR13, MtRR2 and 21 showed similar expression, however paralogs of CaRR13, CaRR15 showed opposite expression (**Figure 6B, 6C and Supplementary Table S4**). Interestingly, only one of the Type-B RRs, CaRR13 showed the neutral selection (Ka/Ks=1) with soybean Type-B RR (**Supplementary Table S4**). Based on all interesting observations we selected CaRR13 as a potential candidate. The hairy-root transformation of *CaRR13* showed its subcellular localization in the nucleus. *In silico* analysis showed that *CaRR13* acts as Myb-like transcription factor which was further validated via interaction analysis of *CaRR13* with NSP2 promoter by Y1H (**Figure 8B**). This further established a link between cytokinin signaling and nodulation as we tried to strengthen our understanding about putative role of *CaRR13*.

**Figure 8.**
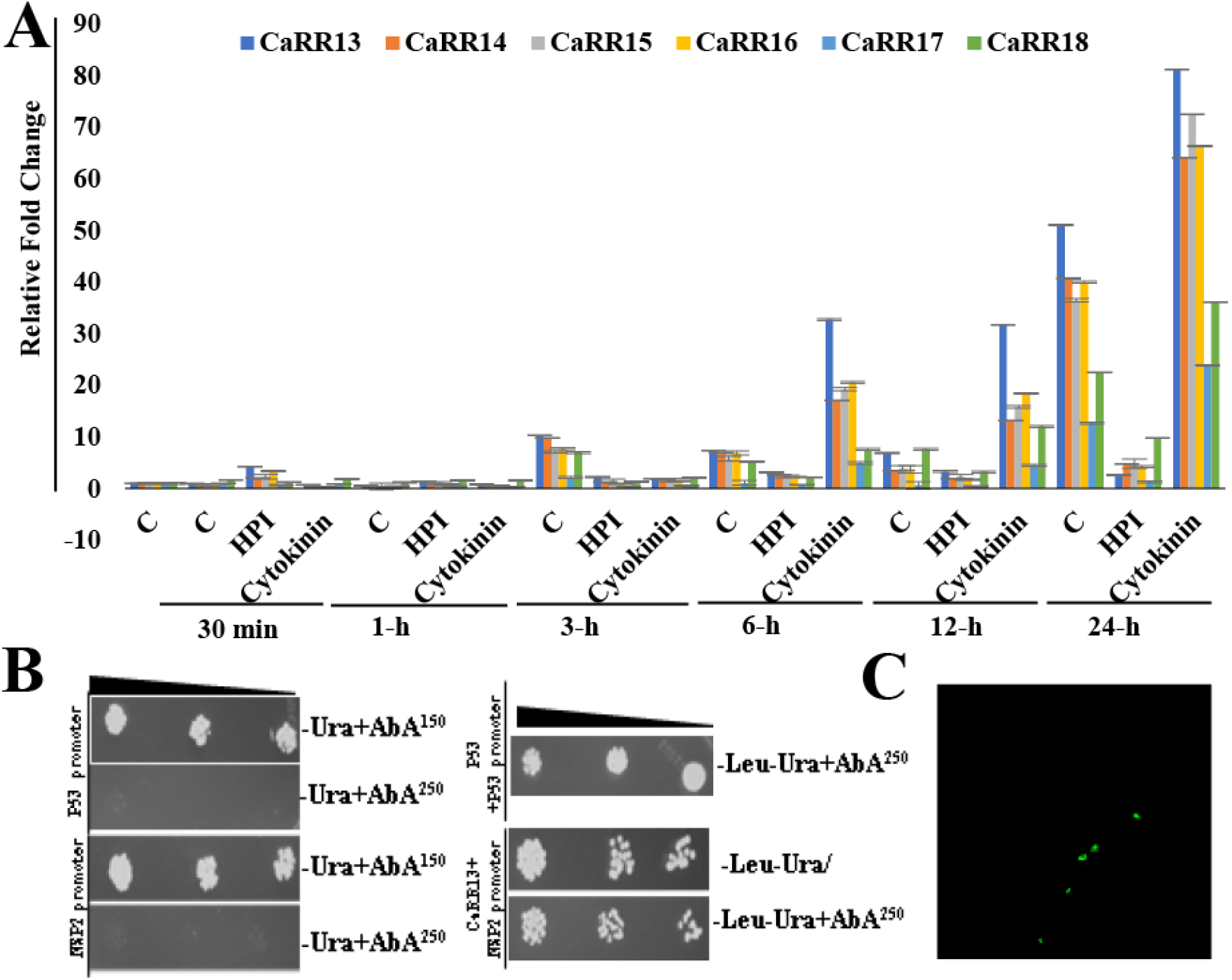
Functional annotation of CaRRs. **A.** Bar diagram represents the expression profile of *Ca*RRs during control (C), *M. ciceri* treatment (expressed as HPI) and cytokinin (2.5X10^−6^M 6-BA). The experiments were carried in triplicates and expressed as means ± S.E. **B.** Interaction of CaRR13 with NSP2 promotor using yeast one hybrid analysis. Strong interaction was observed on Ura and Leu deficient in the presence of aureobasidin A (AbA^150^ and AbA^250^). P53-promoter and AD-Rec-P53 were used as a positive control. **C.** Subcellular localization of CaRR13

## Discussion

Symbiotic interactions of rhizobia with legumes involve perception of flavonoid like substances from root exudates by bacteria and, in turn, bacteria release the Nod factors. Release of Nod factors induces manipulation of phytohormones which is responsible for nodulation. Several cytokinin responsive genes were found to be upregulated during nodulation (Liu et al., 2019; Mortier et al., 2014; Chen et al., 2012). Significant nodulation was observed at 2.5×10^−7^ 6-BA in chickpea even in the absence of rhizobia. However, lower concentration of 6-BA (2.5×10^−5^) reduces the overall growth of plant (**Figure 1A**). Previous investigation showed that few members of the two component signaling (TCS) sensor were active during the cortical cell division also which lead to nodule formation (Held et al., 2014). Therefore, we analyzed the interaction of cytokinin with TCS sensor molecule and how these factors play a putative role in nodule formation. We found the maximum number of TCS members in *Medicago* among the three legumes. The number of TCS members in each organism was in correspondence to the total number of genes in the respective legume species (**Supplementary Table S8**). The higher number of TCS members in *Medicago* compared to chickpea is due to massive local gene duplication in *Medicago* genome (2.63 times more duplication in gene families as compared to soybean) (Young et al., 2011) (**Figure 1B**). The Ks value peaks of orthologous pairs between legumes and *Arabidopsis* was approximately 1.8 revealing the occurrence of duplication ~ 130 Mya. The peaks probably depicted the ‘gamma triplication or whole genome hexaploidy’ commonly shared by all core eudicots (Jain et al., 2013; Varshney et al., 2011; Young et al., 2011).The major peaks of TCS orthologs at ~0.7 among legumes corresponded to a common whole genome duplication (WGD) event (~54-58 Mya) during the origin of papilionoid family. This WGD event coincides with the Cretaceous–Paleogene extinction event which is a hallmark of global extinction event.

Phyloanalysis of HKs revealed different domain containing subclasses. Among the subclasses, chase domain-containing HK (cytokinin receptor, 18 mismatch) were highly evolved across legumes as compared to GAF (ethylene receptor, 6 mismatch) and PHY domain-containing HK (phytochrome receptor, 10 mismatch) (**Supplementary Figure 1B**). Nonetheless, paralogs of cytokinin receptor were also functionally diverged between chickpea and *Medicago* (**Figure 6B and 6C**). Diversification in cytokinin receptors might regulate the signaling components through various mechanisms. This duplication and functional diversification might explain the evolution of HKs involved in bacterial chemotaxis to complex signaling, including phytochrome, ethylene and cytokinin in higher land plants. Interestingly, three cytokinin insensitive HKs (CcHK3, MtHK15 and CaHK3) were highly expressed in nodule and also found to be duplicated among all three legume species (**Figure 6A, 6B and 6C**). These observations suggested that cytokinin independent signaling might also play the regulatory role during nodulation which remained conserve while legume evolution. Few duplicated pairs of TCS members exhibited different expression which depicted its functional divergence through difference in their transcriptional programming (**Figure 6A, 6B and 6C and Supplementary Table 4A and 4B**). All members of CaHPs duplicated from *Medicago* and remain conserved indicating higher similarity. Collinearity analysis of all TCS genes also revealed the closer relationship of chickpea with *Medicago* as compared to pigeonpea. Mostly HPs were segmentally duplicated from soybean to chickpea, *Medicago* and pigeonpea, incidentally, none of CcHPs were duplicated in chickpea or *Medicago*. Continuous elevated expression of the cytokinin positive regulator depicted the importance of cytokinin signaling during the developmental stages of plants especially roots and nodules. This is evident from the fact that cytokinin is produced in root and transported to other places of plant system.

The genome-wide median value of Ka/Ks ratio and higher Ka of RRs depicted that less conservation of RRs as compared to HKs might be responsible for evolution of new functions. Additionally, 2-3 pair of RRs between chickpea and soybean showed Ka/Ks value higher than 1 suggesting the recent positive selection. This might be due to recent WGD in soybean approximately 13 Mya (**Supplementary Table S4**). The comprehensive analysis of duplicated pairs of HKs and RRs in different plants unraveled the divergence of RRs in legume family. Evolutionary analysis suggested that RRs had been diversified in legume-specific lineage to perform various functions in TCS transduction which might help in nodule formation (**Figure 5B, 5C, 5D and 5E**). Few members of Type A and C-RR, were found to be duplicated commonly in all three legumes. The HKs, HPs, Type A and C-RR were found duplicated among all 3 legumes which suggests they have originated from WGD of papilionoid legumes. However, none of the pigeonpea Type-B RRs was found to be segmentally duplicated in chickpea and *Medicago* indicating recent duplication of Type B-RRs during *Medicago* and chickpea evolution (**Supplementary Table S4 and S5**). Through phylogeny and evolutionary analysis, it became evident that Type-B RR diverged between monocot and dicot, and further between legumes and non-legumes depicting the complexity and importance of Type B-RRs in TCS signaling. Therefore, we proposed the involvement of Type-B RRs during root nodulation.

Induction of HKs was observed at 6h of cytokinin treatment and NIN was also following similar expression trend which indicates the connecting link between cytokinin and nodule formation. Various studies showed that nodule-associated transcriptional regulators respond to exogenous cytokinin. Complementation of four HKs confers cytokinin-induced signaling across the kingdom. Previous investigation showed that LHK, LHK1A and LHK3 restored the cytokinin responsive growth, however, LHK2 was unable do so (Held et al., 2014). Response regulators were the main responders of TCS signaling, and most diversified among the TCS components. Novel response regulator candidate, CaRR13 was selected based on its evolutionary importance and transcriptional role in nodule to analyze the mechanistic overview of cytokinin signaling.

## Notes

#### Summary of Updates

supplementary figures and supplementary tables were added

