## Supplementary Figure for "Global analysis of Two Component System (TCS) member of chickpea and other legume genomes implicates its role in enhanced nodulation"

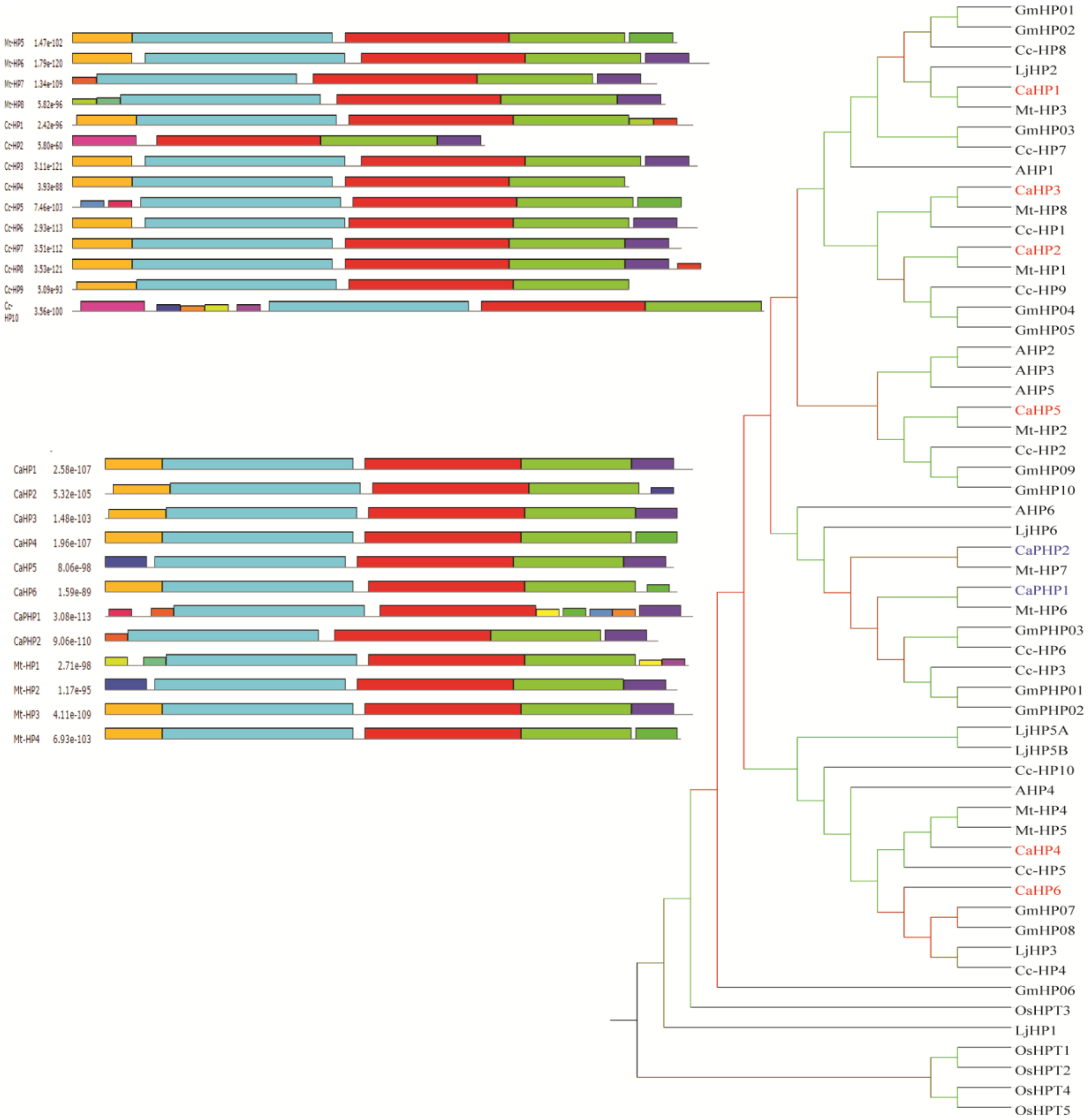


**Supplementary Figure S1.** The phylogenetic tree of histidine phosphotransferase and pseudo histidine phosphotransferase in different crops. The bootstrap values are depicted as color code; green color: maximum and red color: minimum.

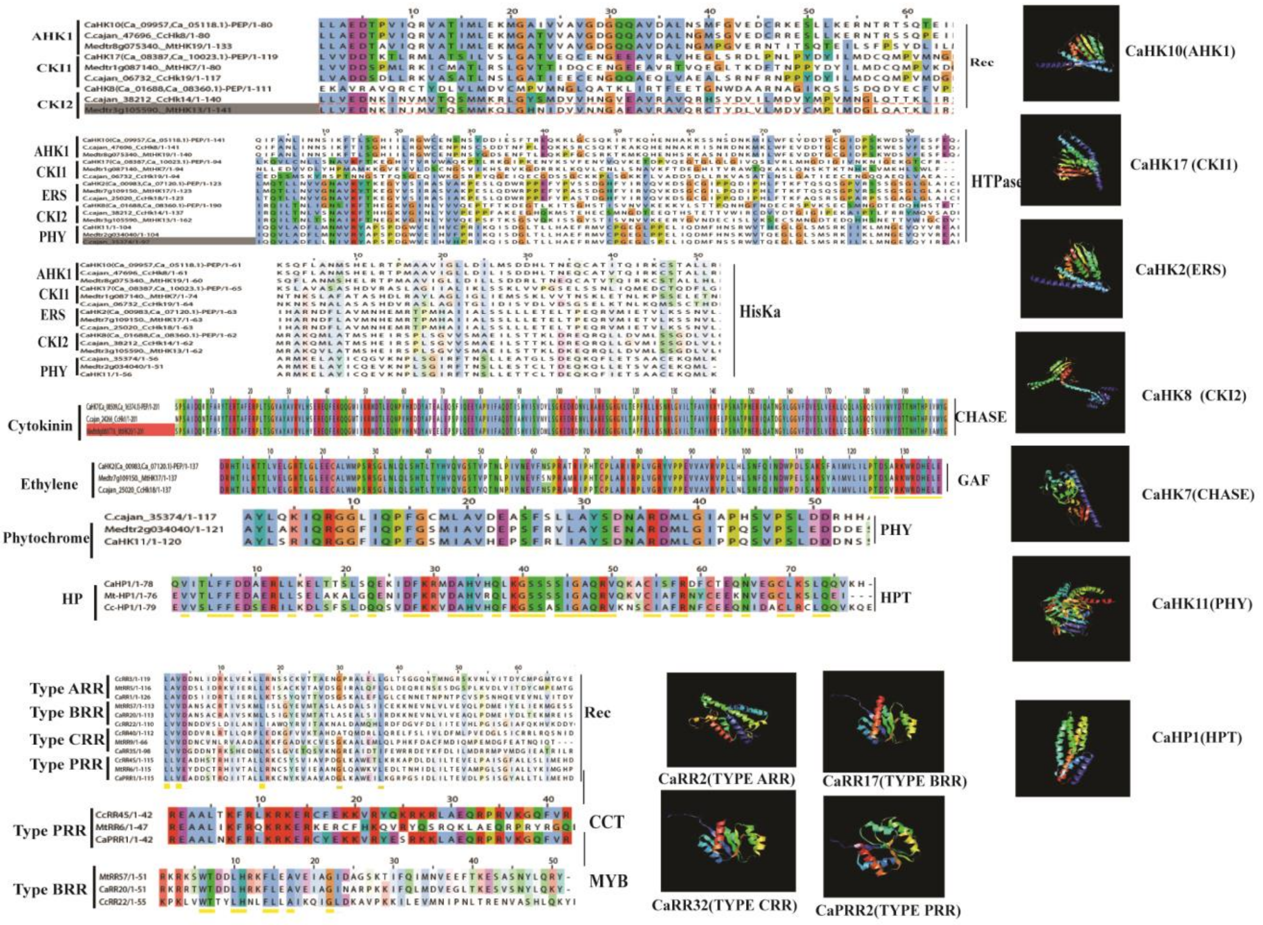


**Supplementary Figure S2.** The multiple alignment of conserved domain and 3-dimensional structure of representative protein from histidine kinase, histidine phosphotransferases and response regulator family.

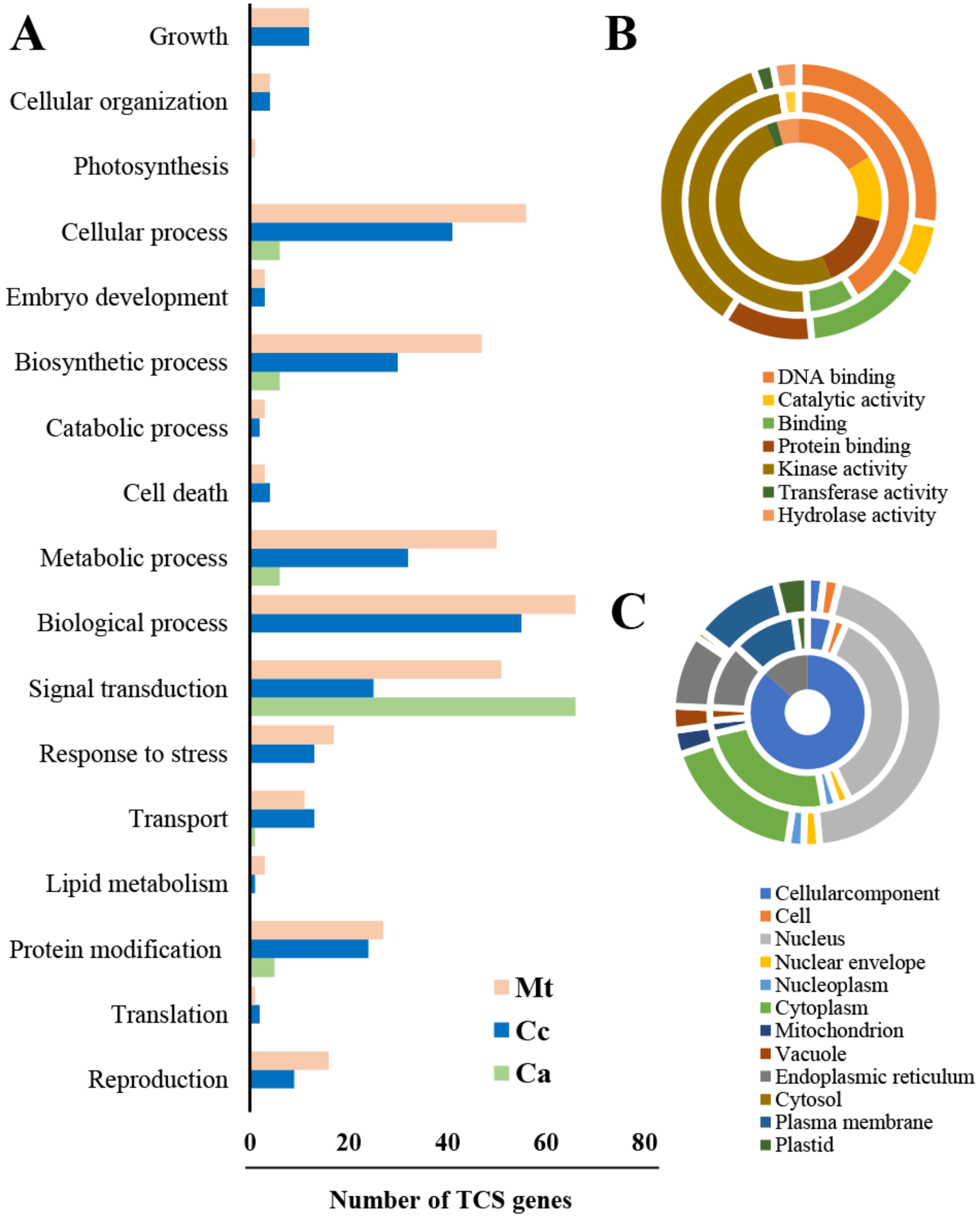


**Supplementary Figure S3** Functional annotation of TCS genes classified in biological process **(A)**, molecular function (B) and cellular component (C) from chickpea (inner circle), *Medicago* (middle circle) and pigeonpea (outer circle).

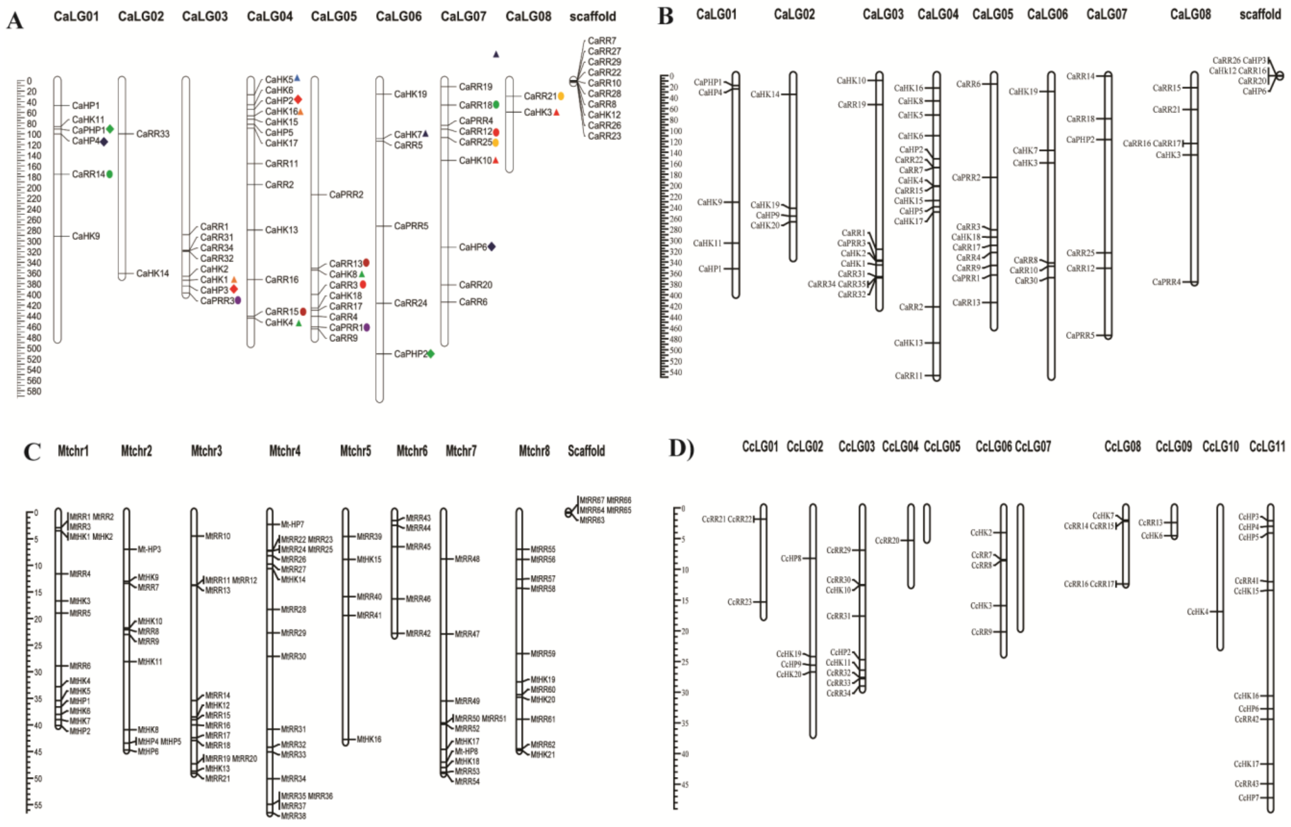


**Supplementary Figure S4.** The chromosome map of TCS genes in kabuli chickpea (A), desi chickpea (B), *Medicago* (C) and pigeonpea (D).

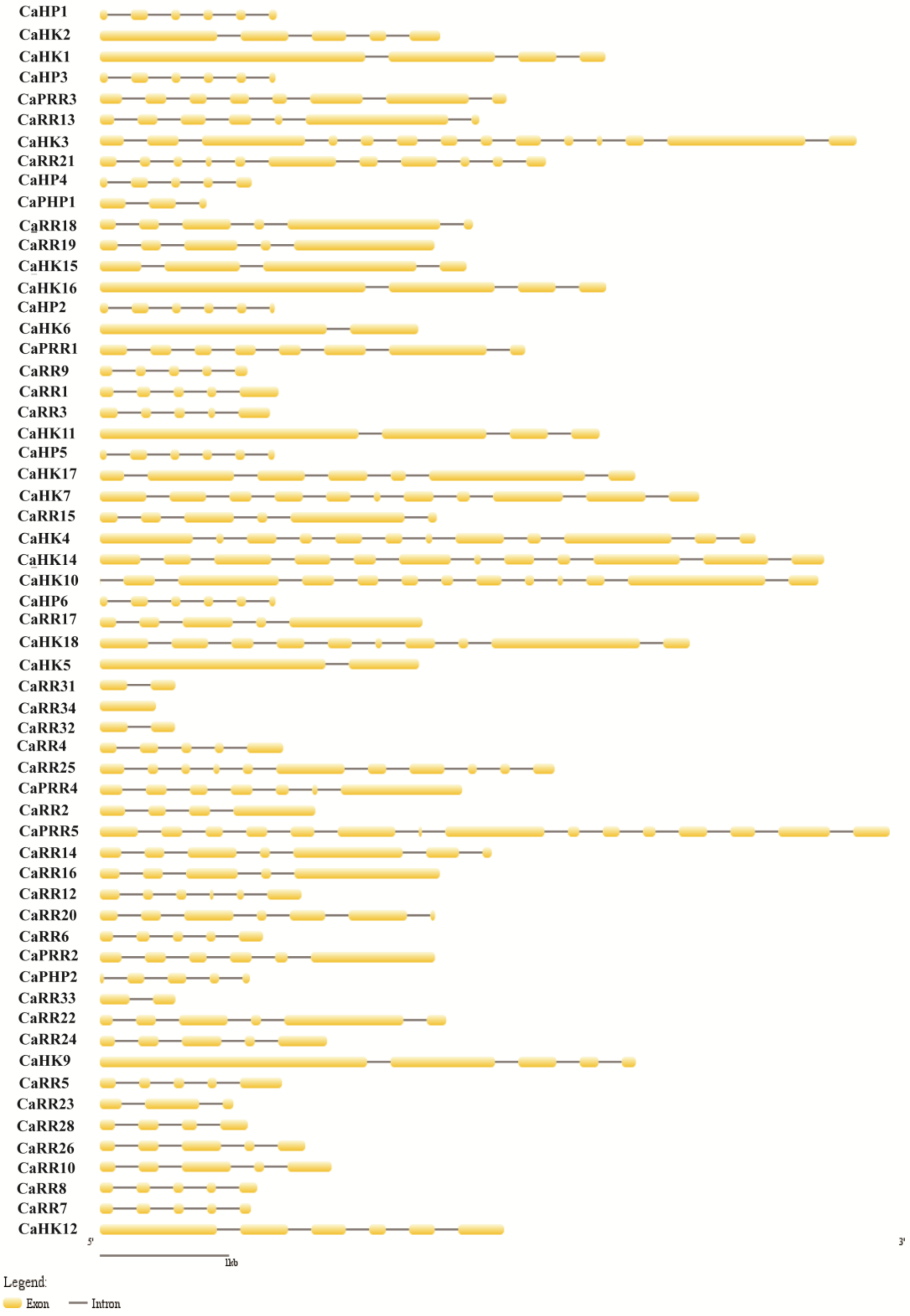


**Supplementary Figure S5.** The GSDS structure of TCS in chickpea

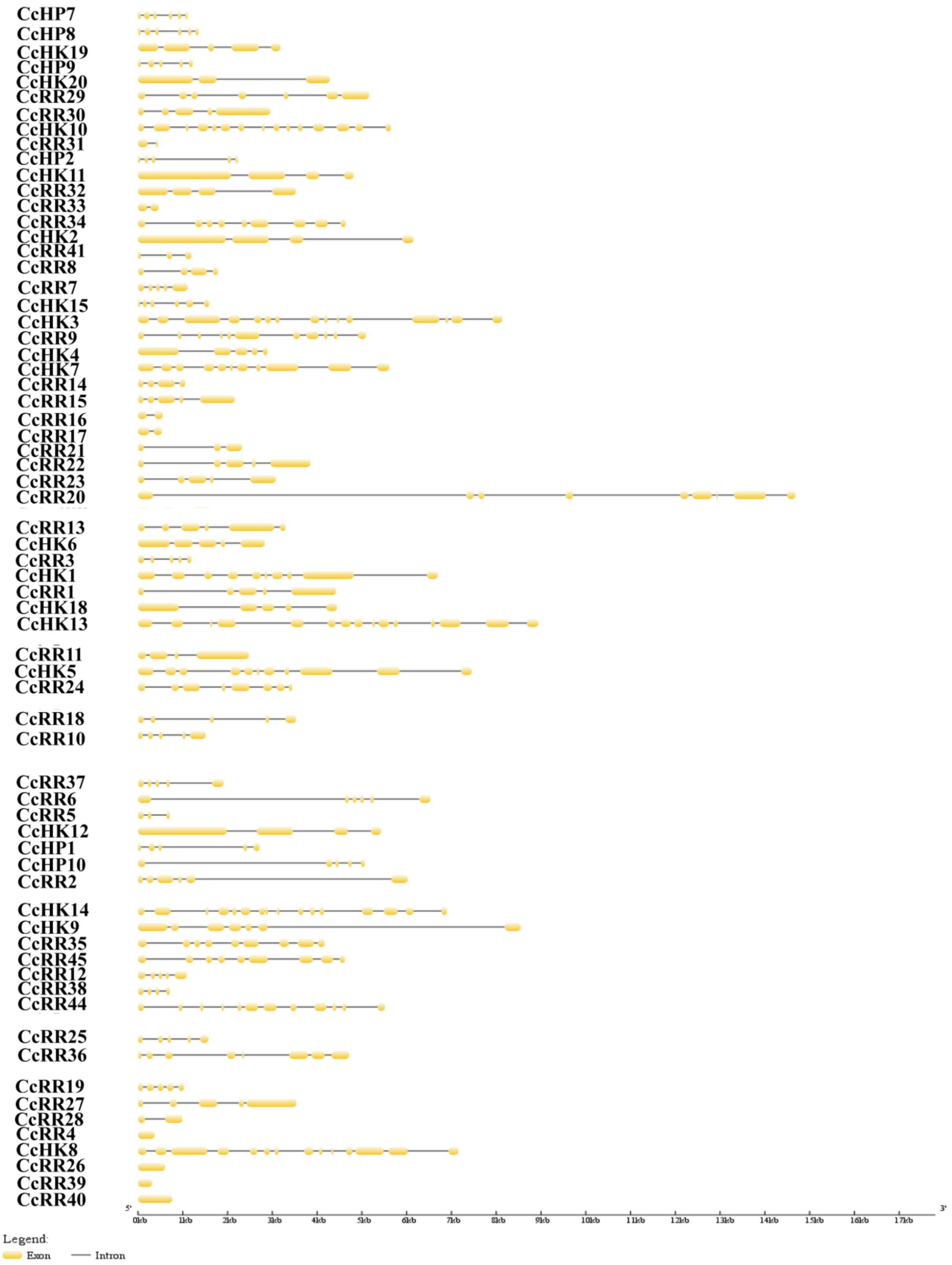


**Supplementary Figure S6.** The GSDS structure of TCS in pigeonpea
